# HiCRes: a computational method to estimate and predict the resolution of HiC libraries

**DOI:** 10.1101/2020.09.22.307967

**Authors:** Claire Marchal, Nivedita Singh, Ximena Corso-Díaz, Anand Swaroop

**Author notes:** Correspondence should be addressed to Claire Marchal or Anand Swaroop.

## Abstract

Three-dimensional (3D) conformation of the chromatin is crucial to stringently regulate gene expression patterns and DNA replication in a cell-type specific manner. HiC is a key technique for measuring 3D chromatin interactions genome wide. Estimating and predicting the resolution of a library is an essential step in any HiC experimental design. Here, we present the mathematical concepts to estimate the resolution of a library and predict whether deeper sequencing would enhance the resolution. We have developed HiCRes, a docker pipeline, by applying these concepts to human and mouse HiC libraries.

## Background

Within mammalian nuclei, chromatin is compacted following a well-defined three-dimensional (3D) organization. Chromosomes remain separated into distinct territories that can be labeled and observed by microscopy [1, 2]. Within each chromosome, the chromatin can be organized into megabase-size domains, called topologically associated domains (TADs) [3, 4]. At the kilobase level, two genomic loci can join together to form chromatin loops [4-6]. This organization is dynamic and changes during distinct stages of a cell’s life including cell cycle [7-9], differentiation [5, 10, 11] and senescence [12, 13]. 3D chromatin organization is associated with gene expression regulation [14-19] and DNA replication timing [7, 20-24], but the relationship between these features is still poorly known. Chromosome Conformation Capture technologies, such as HiC, have permitted access to 3D chromatin interactions genome-wide [25-27] and are among the most common techniques used to explore the relationship between the 3D genome and chromatin associated processes.

HiC libraries are generated by in-nuclei enzymatic digestion of cross-linked chromatin. Digested chromatin is then ligated producing chimeric fragments of neighbor chromatin loci, whichare purified and sequenced pairwise. Interactions between loci, separated by a restriction digestion site, are kept for further analysis [25]. Many laboratories are implementing HiC to examine 3D chromatin interactions to get a better insight into a biological process. “How deep do I need to sequence my HiC library?” is one of the first questions when deciding to perform HiC experiments. The answer is far from being trivial and depends on the chromatin structures to be observed, such as compartments, TADs or loops, as well as the quality of the HiC library [4]. Compartments are very robust across sequencing depths and can be called on small HiC libraries [20, 28]. On the contrary, loops calling requires high resolution HiC obtained by deep sequencing of good quality libraries [6, 28]. High sequencing depth represents a big expense; thus, the first step is usually to sequence the HiC library at low depth (e.g. 100M read pairs), which allows one to evaluate its quality and assess the usefulness of deeper sequencing that is needed for a higher resolution. Accurately predicting the future resolution of a HiC library would allow a user to then choose how deep the library need to be sequenced to obtain a given resolution. While sequencing depth is the main determinant of the resolution, it is important to note that the resolution of HiC data is limited by the restriction enzyme used in the assay [27, 28]. For example, HindIII restriction enzyme produces an average fragment length of 4 kb on the human genome, and thus the best possible resolution will be around 4 kb [26]. For assays using MboI or DpnII, one can achieve a resolution of around 500 bp [26].

A useful definition of the resolution for HiC library was set up by Rao and colleagues [6]. This definition sets the resolution of a HiC experiment as the minimum size window which, when used to calculate the genome coverage, leads to 80% of the windows covered by at least 1000 reads [6, 27]. Mathematically, the resolution is the window size for which 20^th^ percentile of the reads per window equals 1000. This definition is based on the global distribution of the coverage and allows an estimation of the range of interactions that can be observed in a given library, providing an excellent standard for comparison among multiple datasets.

Nevertheless, the relationship between the resolution of a HiC experiment and the size of its library is non-linear [27]. This relationship depends on: 1) the complexity of the library, which can be predicted using published tools such as preseq [29], 2) the percentage of uniquely mapped valid read pairs, directly proportional to the number of de-duplicated reads, and 3) the distribution of uniquely mapped valid read pairs, which can be estimated and predicted by the model presented in this study. We included all these steps within a single pipeline and developed a docker image, called HiCRes. This is the first method to estimate the resolution of a given HiC library and to predict its resolution at a specific sequencing depth.

## Results

### Modeling the HiC fragments distribution on the genome

Elucidation of relationship between the resolution and the quantity of HiC interactions means understanding the relationship between the read coverage distribution, the window size used to calculate the coverage, and the number of read pairs. To model the relationship between these parameters, we used a large high resolution HiC library from human cells GM12878 [6]. We first explored the association of the read coverage distribution to the window size and the number of read pairs. We observed that the distribution of uniquely mapped valid read pairs on the genome varies perfectly linearly with the window’s size used to assess the coverage (Figure 1 A). Similarly, this distribution varies linearly with the number of valid reads (Figure 1 B). Thus, the 20^th^ percentile of coverage can be considered as a linear function of the window size for a given number of reads, as well as a linear function of the number of valid reads for a given window size. From a mathematical point of view, these two functions are partial derivatives of a third function describing the 20^th^ percentile of coverage *versus* the window size and the number of valid read pairs. Here, we show that this last function can be written as Eq. (1), where x is the window size, y the number of valid read pairs and a, b, c and d are some constants to be determined for each library (see methods). Under the hypothesis that the 20^th^ percentile of the coverage only depends on the read number and the window size, Eq. (1) should be sufficient to predict the 20^th^ percentile of the read of coverage given any read number and window size pairs. To confirm this hypothesis, we manually assessed the 20^th^ percentile of the coverage for several read number / window size pairs in the GM12878 HiC library. The function described by Eq (1) perfectly overlaps the observed 20^th^ percentile coverages from different subsamples and window sizes used for the coverage assessment of a high resolution Hi-C from GM12878 cells (Figure 1 C). We realized that Eq (1) accurately describes the relationship between the 20^th^ percentile of the coverage, the read number and the window size. From this equation, the resolution, *i*.*e*. the window size for which the 20^th^ percentile is equal to 1000 reads, can be written as Eq. (2).

**Figure 1.**
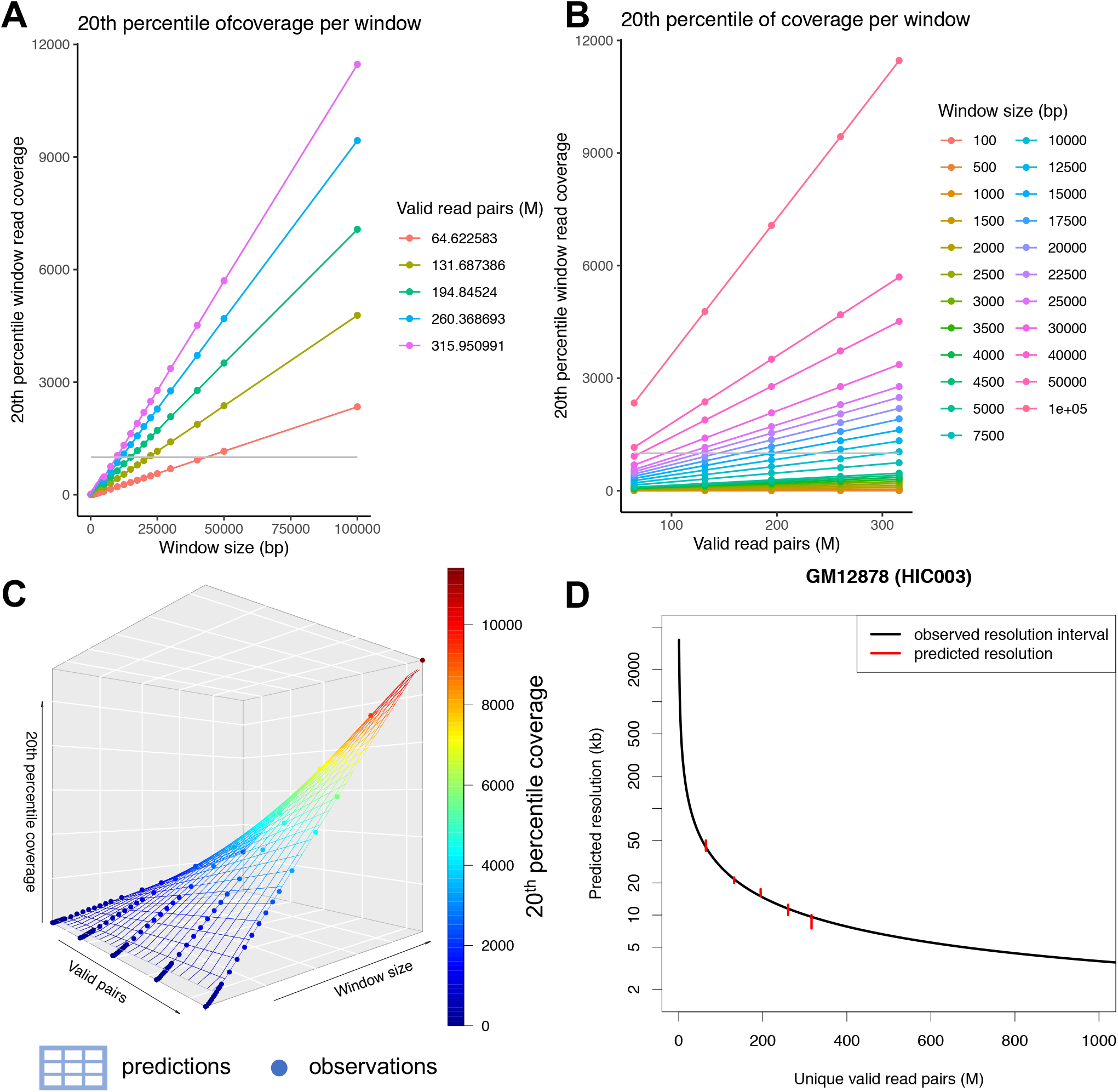
A. Variation of 20^th^ percentile of the coverage with the window size (A) and the subsample size (B) used to calculate the coverage. For each data subsample (A) and each window size (B), this variation is linear. The grey lines represent the limit used to determine the resolution. C. 3D plot showing the prediction of the 20^th^ percentile of window coverage by our model *versus* the window size and the number of valid read pairs. The surface represents the function predicting the 20^th^ percentile for any window size and valid read number, while the dots are the observed 20^th^ percentile for each window size / valid read pairs. The color scale represents the 20^th^ percentile. D. Predicted resolution versus the number of valid read pairs. Predictions are computed using a 100M sequenced read pairs subsample. Observed resolutions of several subsamples are plotted as an interval containing the observed resolution (red segment).

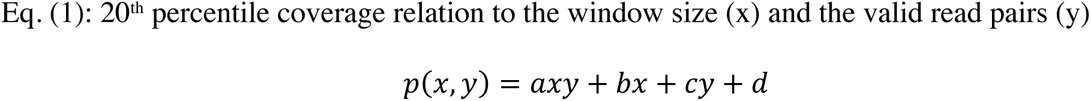

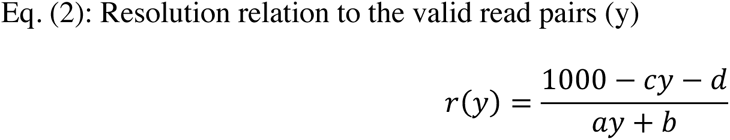

### Validation of the model using published datasets

To validate our model, we used several subsamples of high-resolution HiC datasets that were publicly available (see methods) [6, 30]. We predicted the resolution *versus* the number of valid read pairs using a 100M sequenced read pairs subsample of the library. We then randomly subsampled the GM12878 HiC library into several subsamples. For each subsample, we assessed the number of valid read pairs and measured the interval including the observed resolution (see method). For each subsample, the predicted resolution is within the interval comprising the observed resolution (Figure 1 D). We reproduced this result using two others public HiC libraries, in HMEC and NHEK cell lines (Supplementary Figure 1). For all these three datasets tested, our predictions overlapped perfectly with the observed intervals, thereby validating our model of HiC resolution prediction from the number of valid read pairs.

### Implementation of the pipeline HiCRes

Our model successfully links the number of valid HiC interactions to the resolution. Nevertheless, to predict the sequencing depth required for a library to reach a given resolution, the number of valid interactions needs to be linked to the sequencing depth. To do so, we developed HiCRes, a user-friendly pipeline associating our model to the published tools for measuring any HiC resolution from raw or analyzed data (Figure 2 A). HiCRes is able to predict the resolution *versus* the sequencing depth of any HiC library of 100M read pairs or more. For this purpose, our pipeline measures the library complexity and predicts future yields using the preseq algorithm [29], which we confirmed to be accurate on HiC libraries (Supplementary Figure 2). After estimating the future yield of the library, the percentage of uniquely mapped valid read pairs is evaluated using bowtie2 [31] and HiCUP [32]. Next, the constants a, b, c and d of Eq. (1) are calculated using our model. Finally, the predicted resolution can be calculated for different sequencing depths. For inter-operability, our tool is available as a docker image and can be run on any system where either docker or singularity is installed.

**Figure 2.**
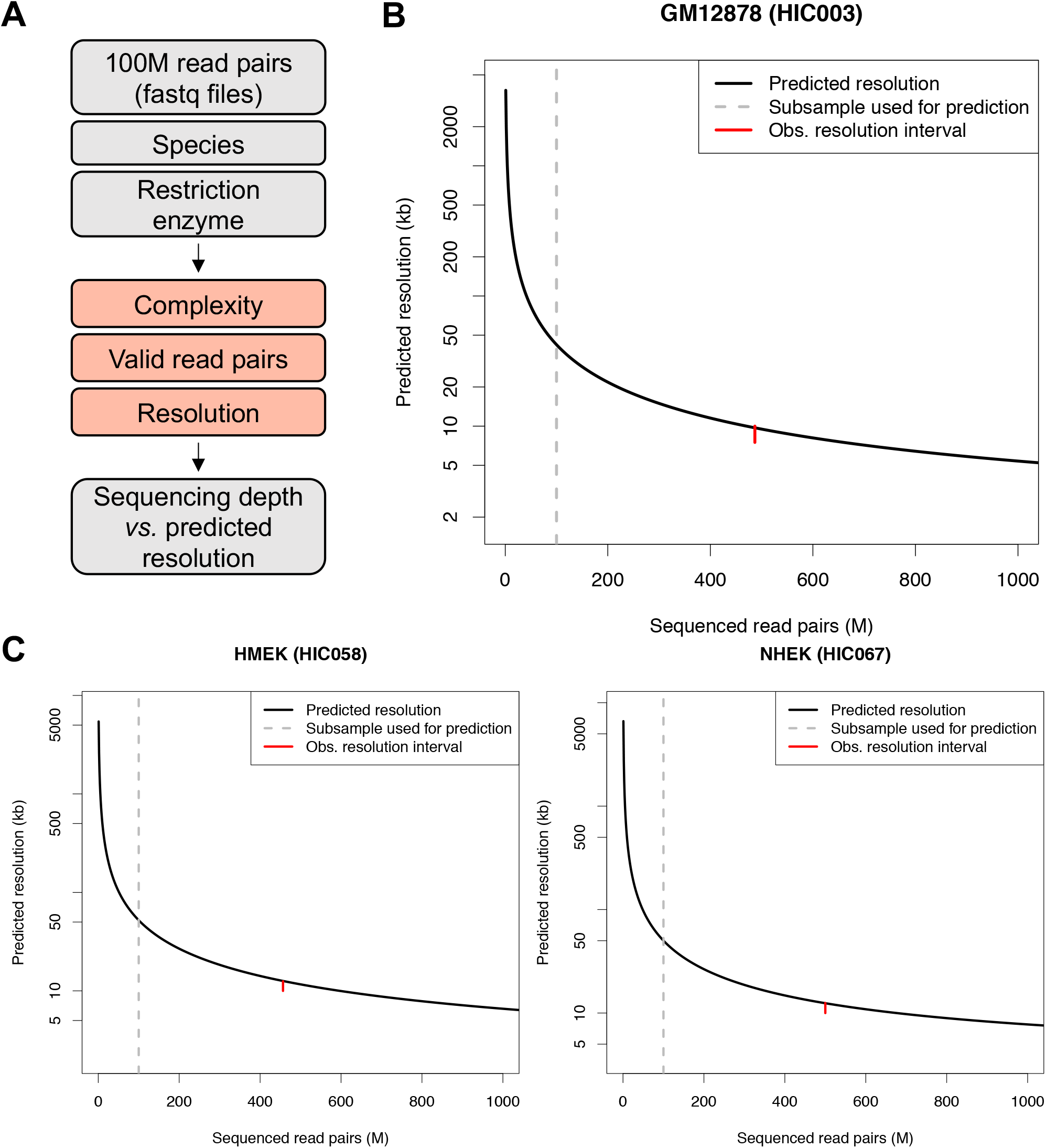
A. The HiCRes pipeline combines preseq which predicts library complexity, bowtie2 and HiCUP which map reads and calculate the percentage of valid read pairs, and HiCRes, which predicts the HiC resolution. This pipeline predicts the resolution of a given HiC library at different sequencing levels. B. Predicted resolution versus the sequencing depth in GM12878 used for the model. Predictions are calculated using a 100M read pairs (grey dotted line) subsample. Observed resolution of the total library is plotted as an interval containing the observed resolution (red segment). C. Predicted resolution versus the sequencing depth in datasets not used for the model: HMEK (left panel) and NHEK (right panel). Predictions are calculated using a 100M read pairs (grey dotted line) subsample. Observed resolutions of the total library are plotted as an interval containing the observed resolution (red segment).

To validate the accuracy of the pipeline to predict the resolution of HiC libraries from raw sequenced reads, we tested HiCRes pipeline on public HiC datasets. We subsampled the GM12878, HMEC and NHEK HiC libraries to 100M sequenced read pairs. We then used HiCRes to predict the resolution each subsampled library will reach for various sequencing depths. To test the accuracy of our predictions, we measured the resolution interval, which is an interval comprising the observed resolution for the whole public library. For each library tested, the prediction of the resolution corresponded with the observed resolution interval (Figure 2 B-D). These analyses confirm the accuracy of HiCRes to predict HiC library resolutions at different sequencing depths based on 100M sequenced read pairs.

### Validation of HiCRes pipeline on diverse HiC conditions

HiCRes has been developed using HiC data from MboI digested chromatin in human cell lines. The use of different restriction enzymes leads to different fragments sizes (Supplementary Figure 3) and could influence the accuracy of our model. To test whether our model and this pipeline can be extended to other species and HiC conditions, we used several public and lab produced datasets. First, we tested HiCRes on HindIII digested chromatin HiC using a public GM12878 HiC library [6] performed using HindIII digestion. We subsampled the library to 100M sequenced read pairs and used HiCRes pipeline to estimate the resolution at various sequencing depths as described above. We then compared the predictions to the observed resolution interval of the full library. The prediction perfectly overlapped with the observed resolution interval (Figure 3 A). Similarly, we tested HiCRes on postmitotic cell types including HiC libraries from MboI digested chromatin of mouse retina [30] (Figure 3 B) and those performed using a kit from Arima technology on purified mouse rod photoreceptors generated in our Laboratory (Figure 3 C). For all these various samples, the predictions perfectly corroborated the observed resolution intervals.

**Figure 3.**
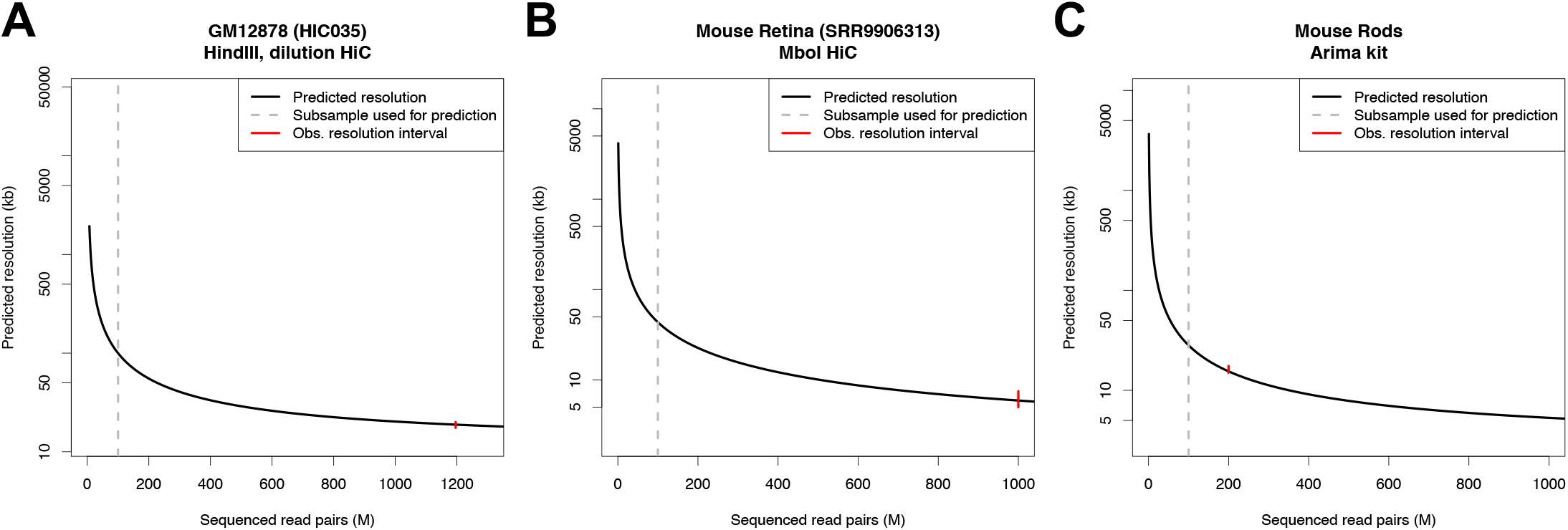
A-C. Predicted resolution versus the sequencing depth in datasets from various species / HiC protocols: GM12878 using HindIII restriction (A), Mouse retina using MboI digestion (B) and Mouse rods using Arima kit (C). Predictions are calculated using a 100M read pairs (grey dotted line) subsample. Observed resolutions of the total library are plotted as an interval containing the observed resolution (red segment).

### Using *cis*-interactions to calculate the resolution

Most tools employed to call 3D chromatin structures use contact maps generated on each chromosome [15, 33]. Thus, only the interactions occurring within the same chromosome, *i*.*e*. the *cis*-interactions, are usually informative for calling compartments, TADs or loops. Accordingly, we added the prediction of the resolution using *cis*-interactions only to our pipeline output. Because *cis*-interactions represent a sub-fraction of all interactions, a lower resolution is expected when using *cis*-interactions only to estimate the resolution, compared to all interactions. For each library tested, our predictions are in accordance with this (Figure 4 A-B, Supplementary Figure 4 A-D). As it would be intuitively expected, we observe a stronger difference between the predictions using *cis* versus all interactions in a library with a high proportion of *trans*-interactions (Figure 4 B), compared to a library with a lower proportion (Figure 4 A).

**Figure 4.**
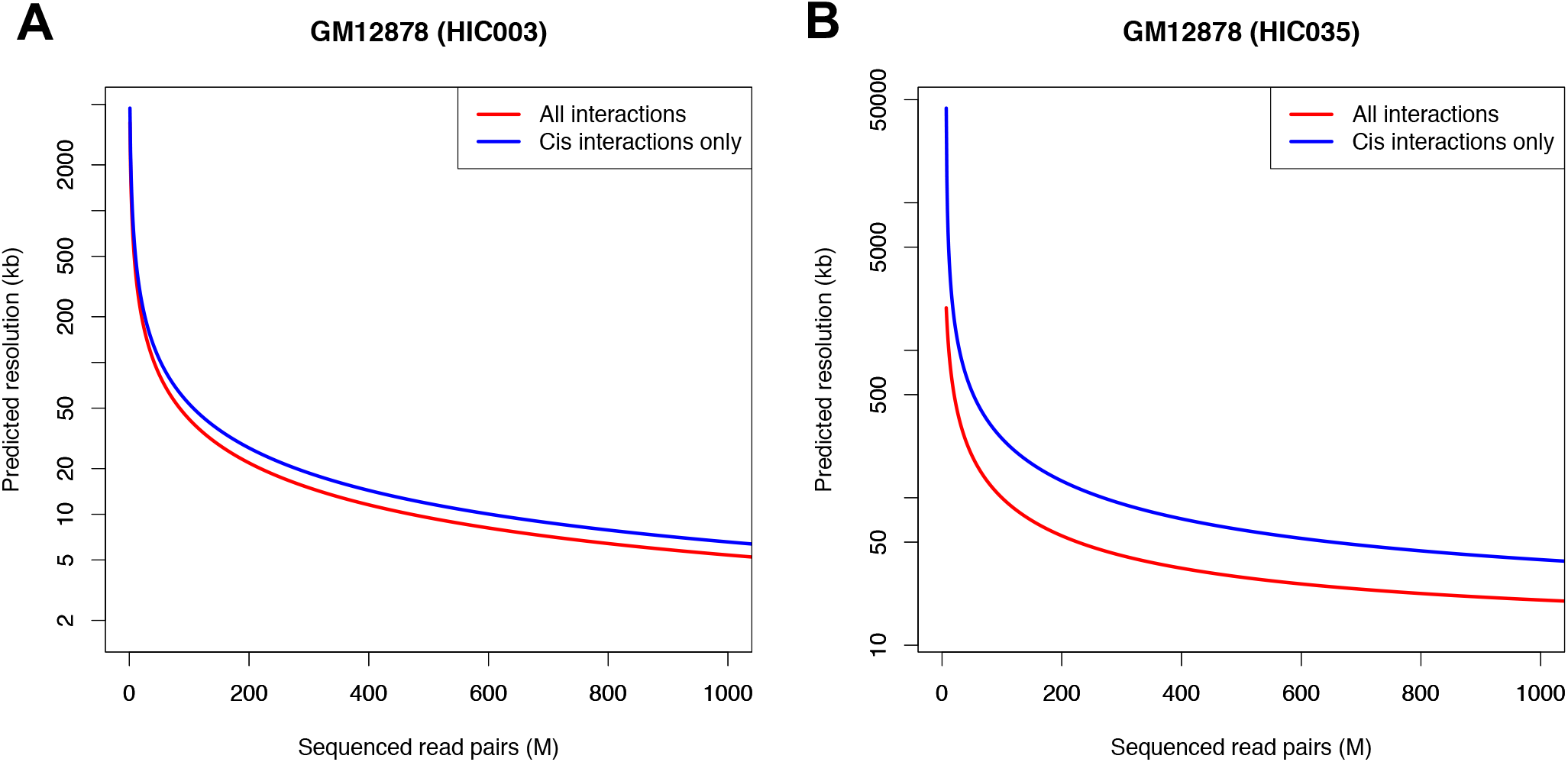
A. Predicted resolution versus the sequencing depth for a HiC datasets in GM12878 with a low proportion of *trans*-interactions (A) and with a higher proportion of *trans*-interactions (B) using all interactions (red) or *cis*-interactions only (blue).

## Discussion

Here, we present HiCRes, a tool to estimate and predict the resolution a given HiC library will reach when sequenced deeper. We demonstrate that HiCRes accurately predicts the resolution of HiC libraries obtained from distinct human and mouse cell types generated using different restriction digestion enzymes. HiCRes is available as a docker image, making it possible to perform different steps of the pipeline using one simple command line.

Two conditions need to be satisfied to apply our model; these are the linear relationships between the 20^th^ percentile of the read coverage with the window size used to calculate the coverage and between the 20^th^ percentile read coverage, and the number of valid interactions. Our pipeline tests whether these two conditions are met and will not produce any estimation or prediction if these conditions are not satisfied. In that scenario, the resolution can be manually measured as described in the method section and by Rao and colleagues [6], but no prediction can be calculated.

HiCRes uses sequenced reads as input to produce a prediction of the resolution *versus* the sequencing depth or already processed HiC data to realize a prediction of the resolution *versus* the number of valid interactions. Using 40 CPUs, HiCRes predicts the resolution of a 200M read pair library in approximatively 5h. Alternatively, processed data can be used as an input for HiCRes. In this case, using 40 CPUs, HiCRes will take approximatively 30 minutes to produce the predictions. Nevertheless, when starting with already analyzed data, the predictions will be done only in relation to the number of valid interactions, not to the library size. Thus, we recommend running HiCRes on raw sequenced reads to predict the resolution that small libraries can reach at deeper sequencing levels. To simply estimate the resolution of a given library (with no need for prediction at different sequencing depths), we recommend running HiCRes directly on processed data.

An important parameter to consider when estimating the resolution of any HiC experiment is the inclusion of *trans*-interactions (*i*.*e*., inter-chromosomal interactions) in the count. The original definition of the resolution by Rao and colleagues included the *trans*-interactions in the valid read count [6]. Nevertheless, in their data, trans-interactions represented around 20% of the valid reads and did not influence drastically the final resolution at high sequencing depths. However, HiC libraries may have a high level of *trans*-interactions when samples (such as tissues) are harder to process. For example, the published mouse retina dataset that we selected possesses a high level of *trans*-interactions [30]. Whether we include these interactions or not in calculating the resolution would have a significant impact on the final result. Given that HiC contact maps are generated usually by chromosome with *cis*-interactions only and are used as input for many tools to perform further analysis (compartments, TADs or loop calling) [15, 33], we recommend using the resolution estimated from the *cis*-interactions only.

The resolution calculated by this approach is a powerful way to estimate the size limit of the chromatin structures to be observed. This prediction can be used to compare different libraries and will help on deciding the sequencing depth needed for a given library. Nevertheless, this number does not directly reflect the quality of the HiC experiment and other quality indicators should be used in complement, such as the proportion of valid interactions, the *cis*-*versus trans*-interactions ratio, the distance-dependent decay of interaction frequency [28] or the reproducibility among replicates [34]. Moreover, the resolution is an average value for the whole genome while local resolutions can be impacted by the read mappability, the presence of restriction digestion sites and the accessibility of such sites to the restriction digestion enzymes. Thus, the measured resolution does not replace the statistical analysis for assessing the local significance of any observed enrichment in a HiC experiment [27].

## Methods

### Datasets used in this study

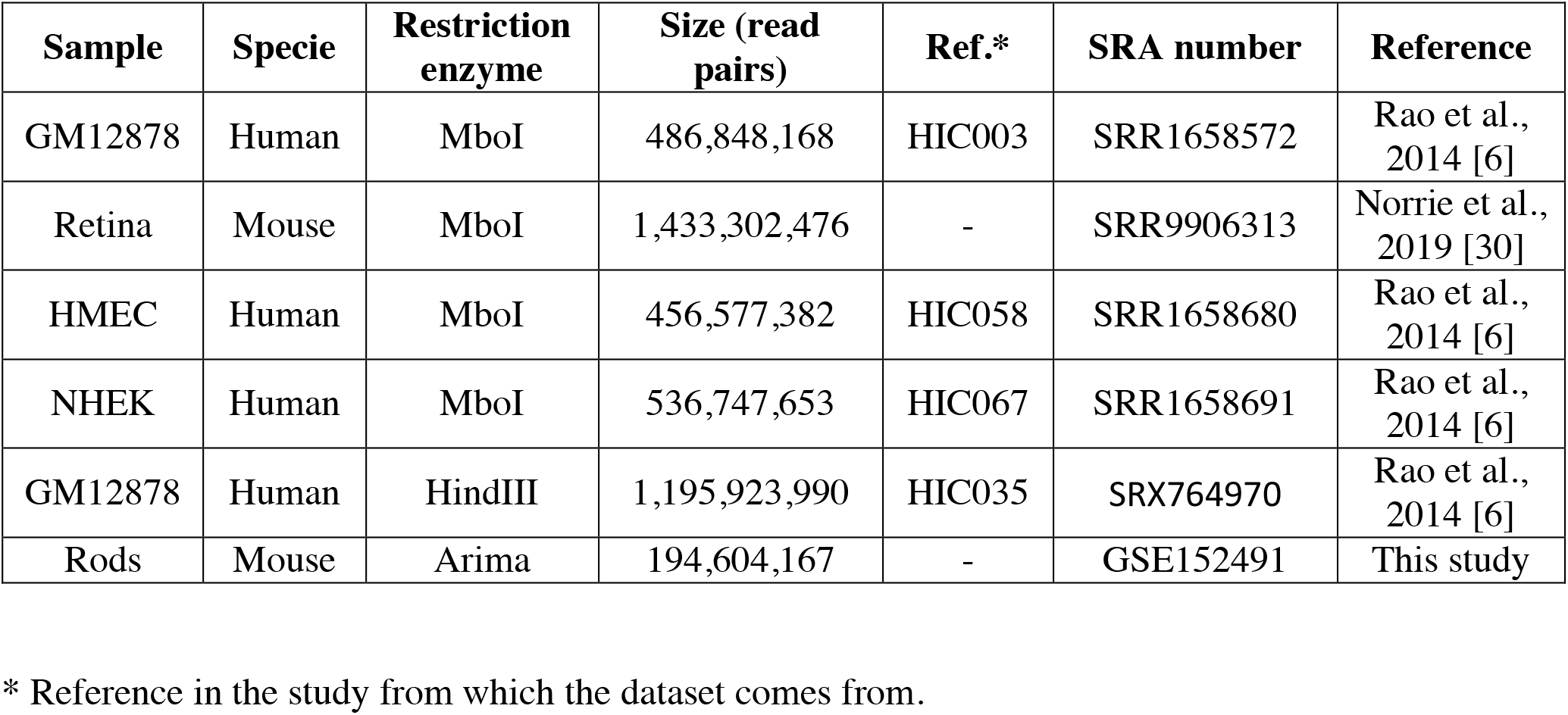

### Subsampling libraries

Libraries are downloaded from SRA and fastq files are extracted using SRAtoolkit (ncbi.github.io/sra-tools/). These files are converted in text files with one complete read pair (sequence and quality) per line. Random lines are then selected using awk bash function and its internal function rand. Seeds for the random extraction are set up as the script running date and time. This method allows the fast extraction of a chosen proportion of reads, while not using the computer RAM. Libraries are subsampled to approximatively 100M, 200M, 300M, 400M and 500M (or the maximum size of the library) read pairs.

### Mapping and filtering

Subsampled libraries are mapped and filtered using bowtie2 [31] and HiCUP [32], on hg38 (human samples) or mm10 (mouse sample), using genomes digested *in silico* by MboI, HindIII or the Arima kit enzymes (Supplementary Figure 3). Proportions of reads pairing, mapping and filtering are calculated with HiCUP. These proportions are constant and independent of the library sequencing depth. Similarly, the proportion of cis-*versus* trans-interaction is independent of the library sequencing depth (data not shown).

### Measuring the observed resolution interval

The final HiCUP output for each subsample is processed through bedtools [35] to calculate the read coverage per window using several window sizes ranging from 100 bp to 100 kb. Then the 20^th^ percentile of the coverage is calculated using R. With these analyses, the 20^th^ percentile of the coverage is measured for each window size. An interval containing the HiC resolution of each subsample is inferred from these values: the minimum of this interval is the larger window size for which the 20^th^ percentile of the coverage is bellow 1000 reads, while the maximum of this interval is the smallest window size for which the 20^th^ percentile of coverage is higher than 1000 reads.

### Combining observed resolution to an equation

The 20^th^ percentile is depending on the number of valid read pairs and on the window size used to calculate the coverage. Thus, it can be written as a function f(x,y) where x is the window size and y the number of valid read pairs. We observed that for a given x value, the 20^th^ percentile is varying linearly with y, which mathematically can be written as Eq. (3).

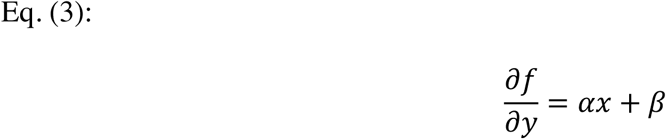

Similarly, for a given y value, the 20^th^ percentile is varying linearly with x, which can be written as Eq. (4).

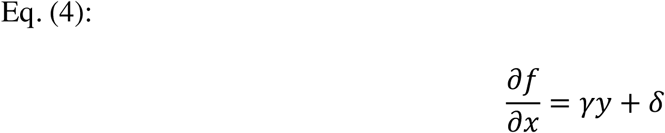

The function f(x,y) satisfying these two equations can be written as Eq. (1).

### Calculating the coefficient of Eq. (1)

The coefficients of Eq. (1) are calculated using a 100M read pairs subsample. The linearity of the distribution of the uniquely mapped valid read pairs with the windows size is controlled using the 20^th^ percentiles from 20 kb, 50 kb and 100kb window size coverage of a 20M valid read pairs subsample. The linearity of the distribution of the uniquely mapped valid read pairs with the read pairs number are controlled by calculating the correlation between 20M, an intermediate number of read pairs and the maximum number of valid read pairs in the sample. Data tested are considered to be linear if the correlation between the 3 points is superior or equal to 0.98. If the linearity is confirmed, 4 distinct datapoints are used to solve the equation for a given library. Here, the measure of the 20^th^ percentile from 20M, an intermediate number of valid read pairs and from 20 kb or 50 kb window sizes are used. Equation (1) is solved using R [36]. From Eq (1), the resolution can be extrapolated as Eq. (2), where x is the number of valid read pairs.

### Estimating and predicting the library complexity

Library complexity is estimated from a 100M read pairs subsample of the library, which will be the minimum size required for the library provided by the user. The library complexity is estimated on the raw mapped read pairs (see mapping and filtering). When both ends of two pairs are mapped on the same position, they are considered as duplicate. Preseq tool is used to predict the library yield at higher sequencing depths from the duplicate distribution. To evaluate the accuracy of preseq on these data, the full GM12878 SRR1658572 library and several subsamples are also analyzed. For each subsample, the non-duplicated reads are counted and compared to the data predicted by preseq based on the 100M read pairs subsample. The perfect overlap between preseq prediction and the observed duplicate rates proves the accuracy of preseq on these data (Supplementary Figure 2).

### Extracting and plotting the resolution *versus* the sequencing depth

Library complexity at several sequencing depths and the associated confidence intervals predicted with preseq are combined with HiCUP statistics to estimate the number of valid read pairs, and valid read pairs in cis for various library sizes. These values and their confidence intervals are then used to calculates the predicted resolution based on Eq. (2).

### HiC on mouse rods

Mouse rods were purified from the Nrl*p*-EGFP C57BL/6J strain as described [37]. All procedures were approved by the Animal Care and Use Committee (NEI-ASP#650). Rods were fixed with 1% formaldehyde for 15 min followed by 5 min incubation with Glycine (125 mM) before cell sorting. Two million purified rods were used for HiC, per instructions from the Arima kit (Arima, # A510008). Libraries were sequenced using Illumina HiSeq 2500.

## Code availability

HiCRes pipeline is available as a docker image on hub.docker.com/r/marchalc/hicres. All the scripts used to produce the figures in this study are available on GitHub, as well as the benchmarking for HiCRes docker (github.com/ClaireMarchal/HiCRes). The dockerfile used to generate the image is also available on GitHub.

## Data accessibility

The HiC dataset on mouse rods generated in this study is available on the GEO database (www.ncbi.nlm.nih.gov/geo/) under the accession number GSE152491.

## Acknowledgments

The authors are grateful to Frederic Mentink-Vigier from the National High Magnetic Field Laboratory (FSU, FL) for providing insights that helped this study. We also thank Linn Gieser and Zachary Batz for assistance with next generation sequencing. This work is supported by Intramural Research Program of the National Eye Institute (ZIAEY000450 and ZIAEY000546) and utilized the computational resources of the NIH HPC Biowulf cluster (http://hpc.nih.gov).

