## Supplemental figures for "HiCRes: a computational method to estimate and predict the resolution of HiC libraries"

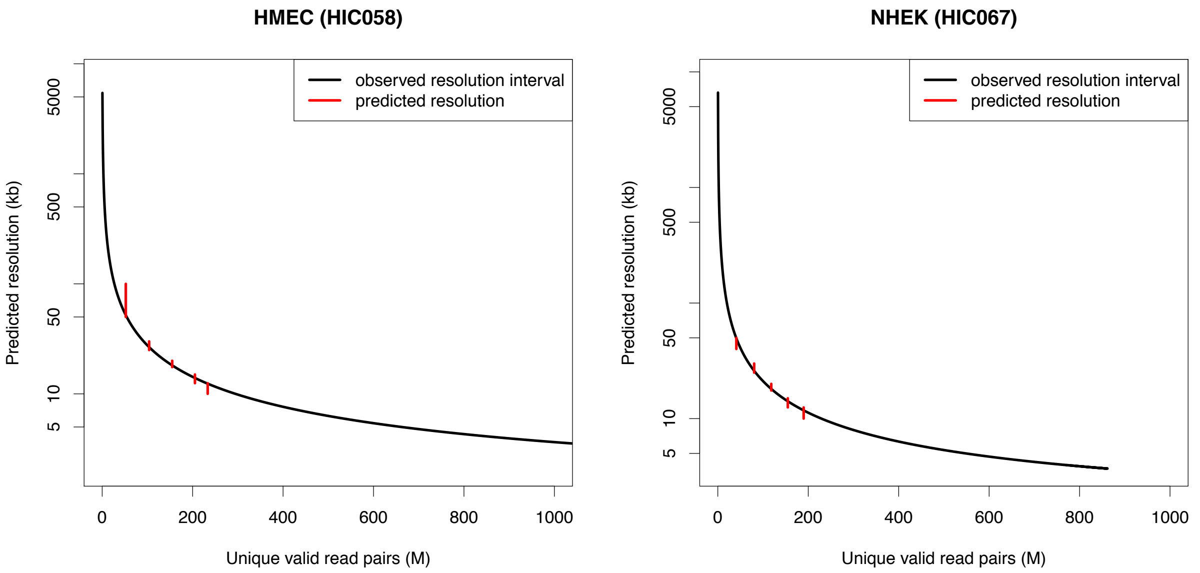


**Supplementary Figure 1.**
Predicted resolution versus the number of valid read pairs in HMEC (left panel) and NHEK (right panel). Predictions are computed using a 100M sequenced read pairs subsample. Observed resolutions of several subsamples are plotted as an interval containing the observed resolution (red segment)

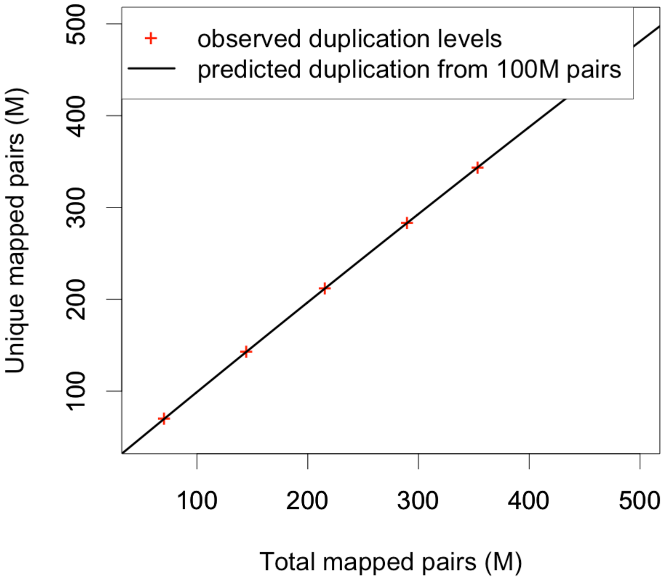


**Supplementary Figure 2.**
Number of total mapped read pairs *versus* unique read pairs as predicted by preseq (black line) using a 100M read pairs subsample or measured in each subsample (red cross).

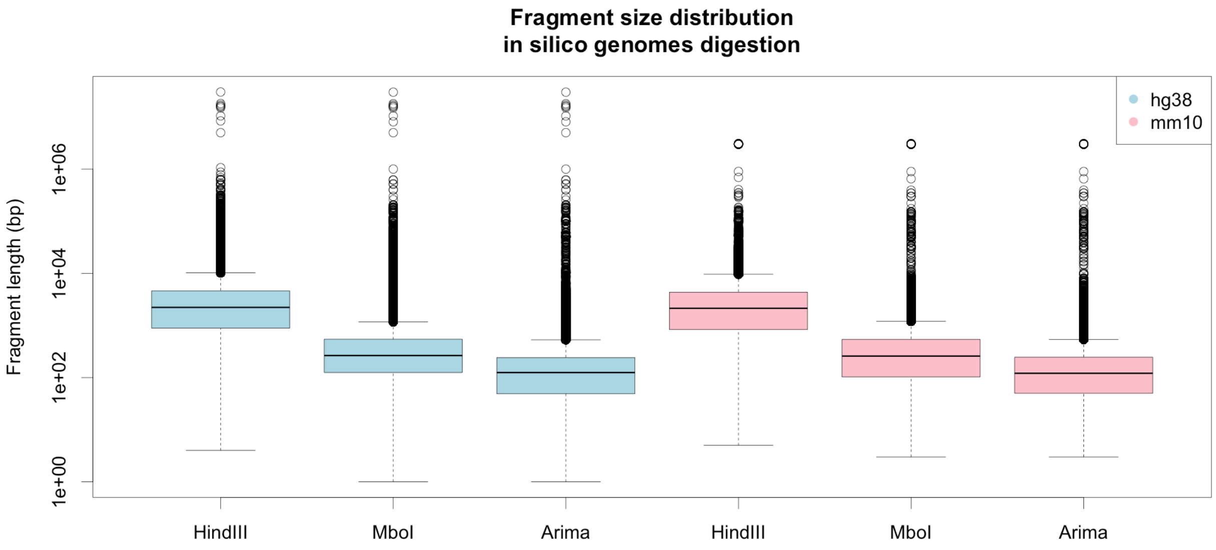


**Supplementary Figure 3.**
*In silico* digestion of hg38 and mm10 genomes for different restriction enzymes.

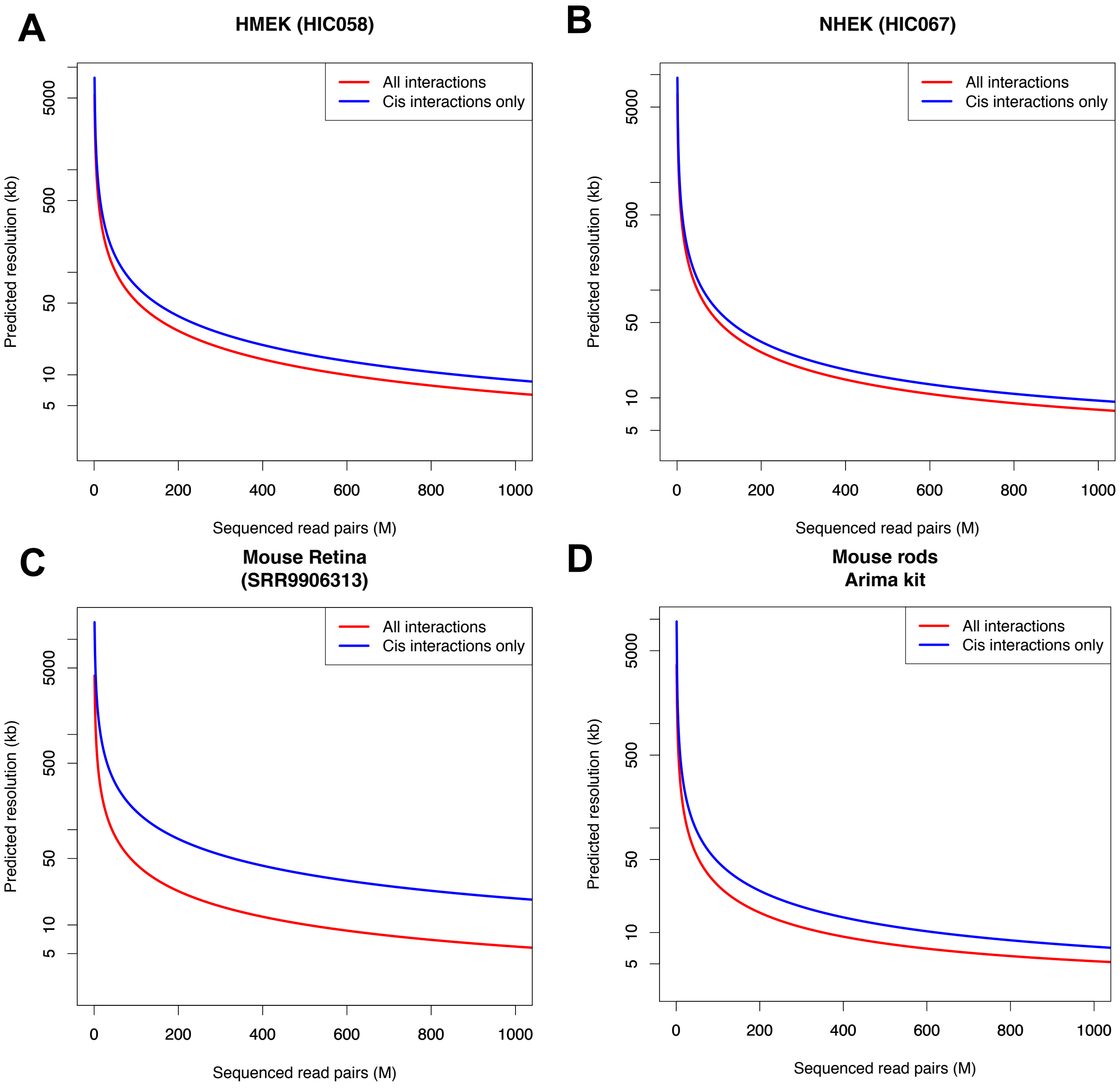


**Supplementary Figure 4.**
A.-D. Predicted resolution versus the sequencing depth in HMEK (A), NHEK (B), mouse retina (C) and mouse rods (D) using all interactions (red) or cis-interactions only (blue).
